# Evolutionary constraints guide AlphaFold2 prediction of alternative conformations and inform rational mutation design

**DOI:** 10.1101/2025.04.08.647782

**Authors:** Valerio Piomponi, Alberto Cazzaniga, Francesca Cuturello

## Abstract

Investigating structural variability is essential for understanding protein biological functions. Although AlphaFold2 accurately predicts static structures, it fails to capture the full spectrum of functional states. Recent methods have used AlphaFold2 to generate diverse structural ensembles, but they offer limited interpretability and overlook the evolutionary signals underlying predictions. In this work, we enhance the generation of conformational ensembles and identify sequence patterns that influence alternative fold predictions for several protein families. Building on prior research that clustered Multiple Sequence Alignments to predict fold-switching states, we introduce a refined clustering strategy that integrates protein language model representations with hierarchical clustering, overcoming limitations of density-based methods. Our strategy effectively identifies high-confidence alternative conformations and generates abundant sequence ensembles, providing a robust framework for applying Direct Coupling Analysis (DCA). Through DCA, we uncover key coevolutionary signals within the clustered alignments, leveraging them to design mutations that stabilize specific conformations, which we validate using alchemical free energy calculations from molecular dynamics. Notably, our method extends beyond fold-switching, effectively capturing a variety of conformational changes.

## 1 Introduction

The three-dimensional structure of proteins is crucial to their function in biological processes, yet predicting the diverse conformations that proteins can adopt is a significant challenge in structural biology. Recently, AlphaFold2 (AF2) [1], a deep learning model for protein structure prediction, achieved unprecedented accuracy and marked a major breakthrough in computational structural biology. Nevertheless, despite its remarkable performance, AF2 is primarily designed to predict a single dominant conformation, and its ability to capture multiple functional states of a protein remains limited [2, 3]. Several strategies have been developed to improve conformational sampling using the AlphaFold2 algorithm. However, these methods often lack interpretability, particularly in terms of the evolutionary signals that drive the conformational transitions. As a result, the connection between sequence divergence, evolutionary pressures, and the resulting structural states remains poorly understood in many cases. Existing strategies for sampling protein conformations with Alphafold2 comprise two main approaches. The first class of methods involves incorporating conformational information into either the training data or the framework itself, while the second concerns manipulating the Multiple Sequence Alignment (MSA) input. Within the first category, AlphaFlow [4] fine-tunes AF2 by introducing a flow matching objective, which generates structural ensembles by training with both experimental data and molecular dynamics simulations. C-Fold [5] retrains AlphaFold2 removing the template track from the architecture and using a conformational split of the Protein Data Bank (PDB), which enables the model to learn structure flexibility from experimental data. Another method [6] integrates state-annotated structure databases into AlphaFold2’s structural templates, specifically for G-protein-coupled receptors. The second class of approaches involves manipulating the input of AlphaFold2. One of the most successful techniques involves reducing the depth of the MSA by randomly sub-sampling sequences [7], which can increase uncertainty and enhance structural diversification when the number of retained sequences is very low. Similarly, another method [8] also sub-samples the MSA but enhances model diversity by adjusting the number of non-center sequences excluded during AlphaFold2’s initial clustering step, thereby modulating co-evolutionary signals. An alternative strategy involves masking columns of the MSA similarly to SPEACH AF method [9], altering the information within the alignment and forcing AF2 to reconstruct the masked region to explore alternative conformations. Furthermore, AF-Cluster [10] generates diverse protein conformations by clustering aligned sequences based on similarity. Using these clusters as separate inputs for AF2 effectively leads to sample distinct conformational states of fold-switching proteins.

We embrace this last perspective and hypothesize that AF2 predictions are biased toward conformations with prominent coevolutionary signals in the MSA, rather than necessarily favoring the most thermodynamically stable state. This effect stems from the interplay between competing coevolutionary signals within the alignment, which correspond to distinct structural states and shape the predicted conformation. Building on this hypothesis, our aim is to detect metastable states with a limited number of predictions, prioritizing efficiency and interpretability. The small cluster sizes associated with alternative states from existing methods make it challenging to identify evolutionary signatures linked to contacts from different conformations. By generating more comprehensive groups, we can perform reliable statistical analysis of the clustered sequences and identify key regions within the alignments that influence conformational changes.

We combine agglomerative clustering with representations generated with a protein language model and demonstrate superior detection of high-confidence alternative states compared to density-based clustering on a set of fold-switching proteins. Clustering a large number of sequences allows us to apply Direct Coupling Analysis (DCA) [11–14], a statistical framework which models co-evolutionary signals within sequence alignments to infer direct residue interactions, and identify co-evolved pairs that are crucial for the alternative prediction. Focusing on co-varying residues that interact specifically in the alternative state, we design targeted mutations that preferentially stabilize particular conformations by measuring frequency shifts with respect to the full alignments. We also validate a specific mutation pair stabilizing an alternative fold using alchemical molecular dynamics (MD) simulations [15]. Figure 1 shows a schematic overvew of the pipeline. We showcase the versatility of the developed strategy by applying it to various conformational changes beyond fold-switching. Overall, by uncovering evolutionary signals that underlie structural transitions, we distill essential sequence-structure relationships while minimizing computational overhead.

**Figure 1:**
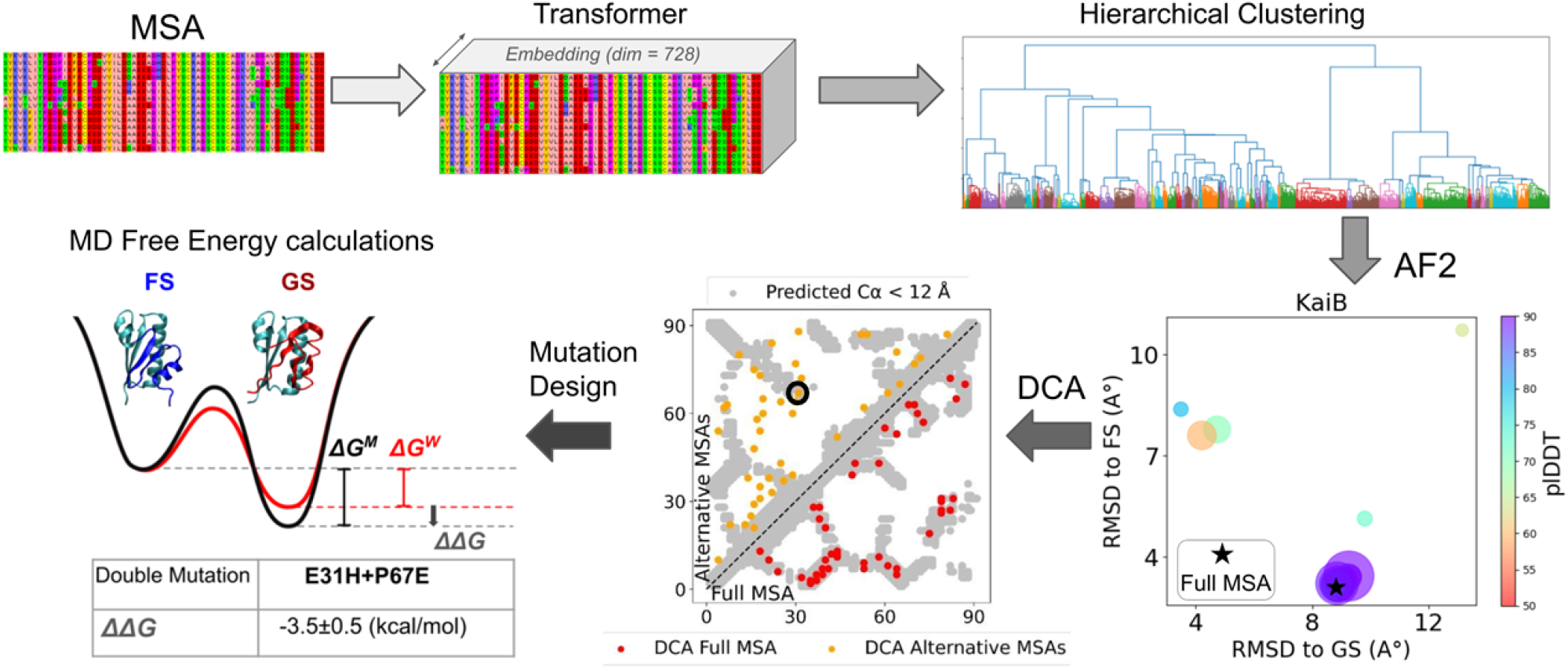
Overview of the pipeline, illustrated with the KaiB fold-switching protein. The aligned homologous sequences are embedded using the MSA Transformer and clustered to input AF2. Predictions either match the default state (FS for KaiB) or the alternative state (GS for KaiB). DCA of alternative clusters helps identify mutated pairs for stabilizing the alternative state, as validated through MD-based free energy calculations.

## 2 Clustering the MSA

### 2.1 Method

The pioneering study [10] introduces clustering of aligned sequences as input for AlphaFold2 to predict diverse conformations, using DBSCAN [16]. The DBSCAN algorithm identifies dense regions, where sequences are closely packed, and separates them from low-density areas relying on two critical hyperparameters: the maximum distance between points in a cluster (*ϵ*) and the minimum sample size, which is the smallest number of points required to form a dense region. Through a grid search, the *ϵ* value is optimized to produce the highest number of clusters while maintaining a very low minimum sample size, to ensure high heterogeneity of sequence groups. However, this approach results in a highly fragmented clustering landscape characterized by numerous small clusters, with only a few resulting in meaningful alternative predictions. Many clusters produce low-confidence structures that significantly deviate from both the ground and the fold-switched conformation. Since the method requires running AlphaFold2 on each group, it is to notice that such abundance of small clusters significantly slows down the pipeline and limits scalability to large datasets. Furthermore, the presence of a “halo” of data points that fail to meet the density criterion leads to significant information loss: biases in sequencing and recombination events can create discontinuities in the sequence landscape, causing density-based approaches to misclassify many sequences as noise.

We develop an approach focused on detecting metastable states with a limited number of predictions, enabling larger and more cohesive clusters that facilitate the identification of critical sequence features. We propose an agglomerative hierarchical clustering (AHC) approach [17], using Ward’s linkage method to merge clusters. This method iteratively merges clusters while minimizing the increase in total within-cluster variance. We iteratively decrease the method’s maximum cluster number parameter until each cluster contains at least 20 elements, to prevent the creation of small clusters that might not be representative. The performance of DBSCAN are here re-evaluated choosing an increased minimum sample size of 20 (with *ϵ* set as previously illustrated). Finally, we introduce the clustering of representations from the last hidden layer of the MSA Transformer model [18], a deep learning architecture designed to extract rich, context-aware embeddings from sequence alignments of protein families. Leveraging the attention mechanisms to integrate information across homologous sequences in an alignment, the MSA Transformer captures both residue-level dependencies and evolutionary constraints. We choose to use these representations to transition from a sequence-based similarity measure to a structured continuous latent space, while preserving evolutionary information. AlphaFold2 predictions are performed using ColabFold [19], following a protocol with three recycling iterations, generating five models per prediction, and without employing any structural templates. We report the results for the top-ranked model, as ranked by the pLDDT score.

### 2.2 Results on Fold-Switching Proteins

The AHC clustering method consistently identifies a larger fraction of sequences associated with alternative predicted conformations compared to DBSCAN (Figure 2). This improvement is crucial for downstream analyses, as it favors the identification of critical signals within the MSA that modulate the switching between states. AHC achieves a slight improvement over direct sequence-based clustering by incorporating representations from the MSA Transformer [18]. However, the overall performance difference between these two approaches remains relatively small, suggesting that sequence similarity alone is often sufficient to guide effective clustering, while evolutionary-based embeddings provide only a marginal refinement.

**Figure 2:**
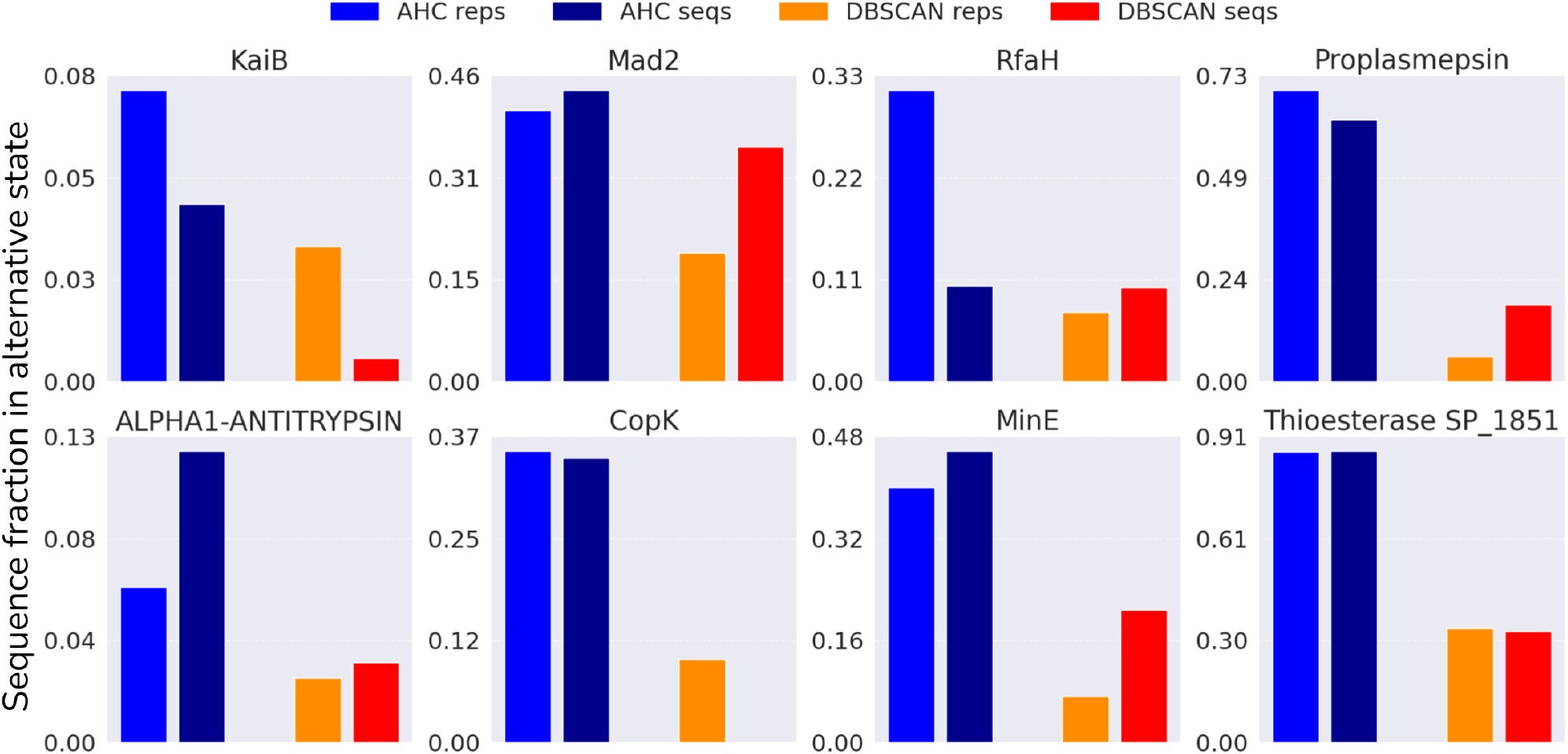
Fraction of sequences associated with alternative state predictions, relative to the full MSA depth, for different clustering methods (AHC and DBSCAN) and inputs (sequences and MSA Transformer embeddings).

Based on these findings, we use MSA Transformer-based AHC clustering for the subsequent analysis. Figure 3 presents the RMSD of cluster-based AlphaFold2 predictions with respect to experimental PDB structures of the two distinct folded states across eight fold-switching proteins. This analysis demonstrates that the clustering strategy produces a well-balanced distribution of conformations, effectively capturing structural variability across different folds. The number of false positives—defined as predictions that deviate significantly from both experimental reference structures—is minimal compared to the foundational work, highlighting the improved reliability of our approach. Moreover, the predicted structures exhibit consistently high pLDDT scores, indicating AlphaFold2’s strong confidence in their accuracy. These results underscore the effectiveness of the clustering strategy in capturing meaningful conformational diversity while minimizing noise.

**Figure 3:**
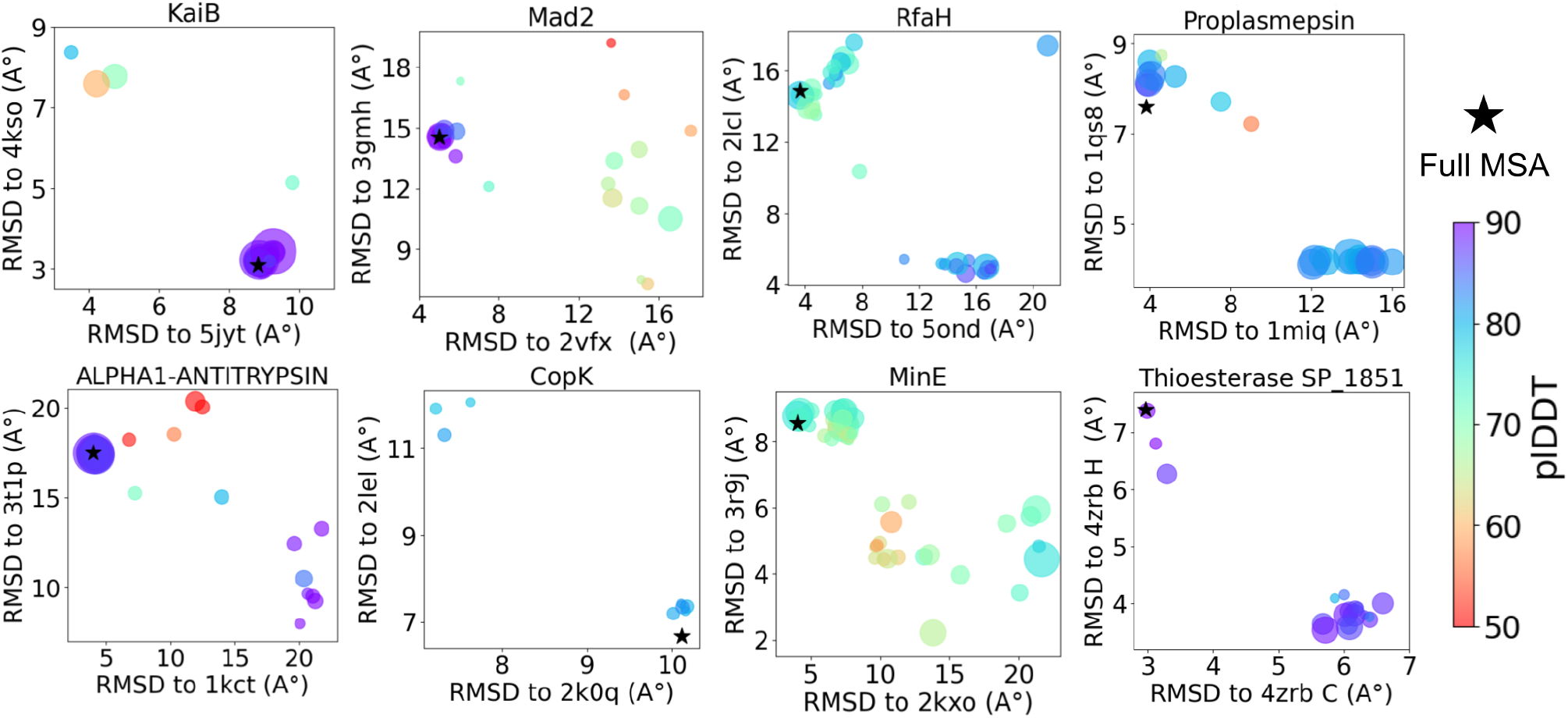
AlphaFold2 structure prediction with clustered MSA for eight fold-switching proteins: RMSDs are computed with respect to experimental PDB structures. The size of circles reflects the cluster size.

To further assess the robustness of the method we applied the clustering strategy to two additional fold-switching proteins—CLIC1, and lymphotactin—which the foundational study had previously failed to resolve into alternative states (Figure 4). The third protein mentioned in the study is Selecase, which poses a particularly challenging case due to the limited number of sequences in its full MSA (only 140). As the method relies on sufficiently populated alignments, it is not applicable to this protein. For CLIC1, we fail to identify a cluster corresponding to the alternative conformation. While some clusters exhibit high RMSD relative to the default AlphaFold2 prediction, the observed structural variability does not align with a meaningful fold-switching event. In contrast, for lymphotactin, we succeed where the original approach failed, identifying a cluster of 90 sequences associated with the alternative folded state. This result highlights the improved effectiveness of our clustering strategy in capturing fold-switching events by leveraging structural signals in the MSA when a sufficient number of sequences is available.

**Figure 4:**
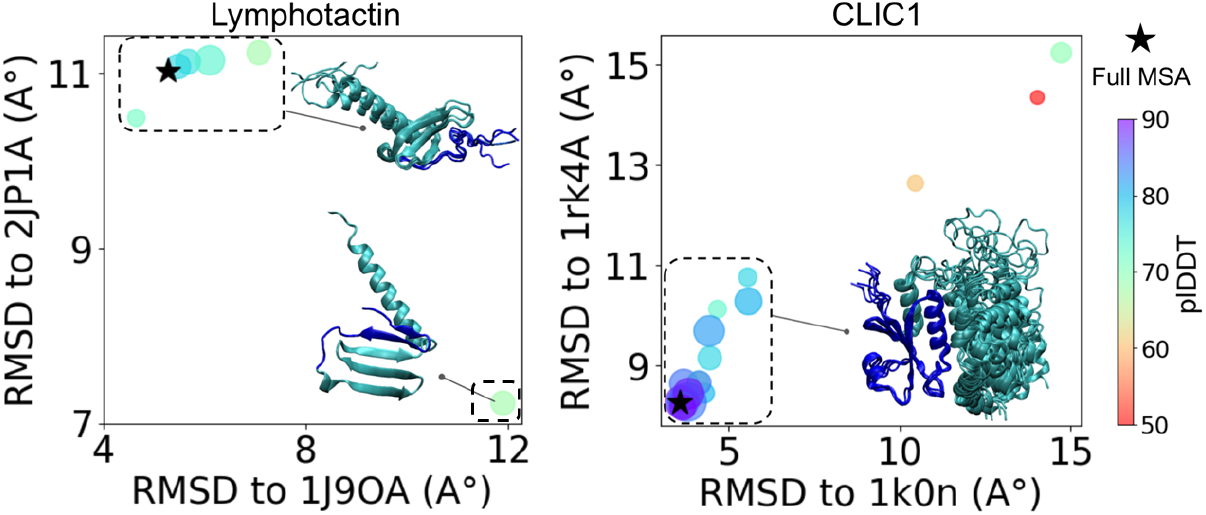
AlphaFold2 structure prediction with clustered MSA for Lymphotactin and CLIC1: RMSDs are computed with respect to experimental PDB structures. The size of circles reflects the cluster size.

We also perform Principal Component Analysis (PCA) on representations from the Evoformer block, which is a crucial component of AlphaFold2’s architecture, to explore how structural variations emerge at different stages of its workflow. The Evoformer block consists of two main modules: the MSA Transformer module, which encodes evolutionary relationships between sequences, and the pair representation module, which refines residue-residue interactions using triangle attention to update structural parameters. These modules continuously exchange information, enabling the model to integrate co-evolutionary constraints and enhance the accuracy of structural predictions. In most systems, the first two principal components of the single representations provide limited separation between the two states (Figure S2 in SI). In contrast, PCA of the pair representations (Figure 5) reveals a clear distinction between the two conformations, underscoring the importance of incorporating MSA-derived information into residue-residue interactions. This pattern is further supported by the cosine distance distribution of pairwise representations (Figures S3 in SI), which shows high intra-state similarity while maintaining a clear separation between the default and alternative conformations. These findings indicate that AlphaFold2 captures fold-switching events at the abstract representation level. The observed separation suggests that both structural and evolutionary constraints are encoded within the Evoformer stage, effectively embedding multiple conformations in its internal representations before the structural module refines the final prediction.

**Figure 5:**
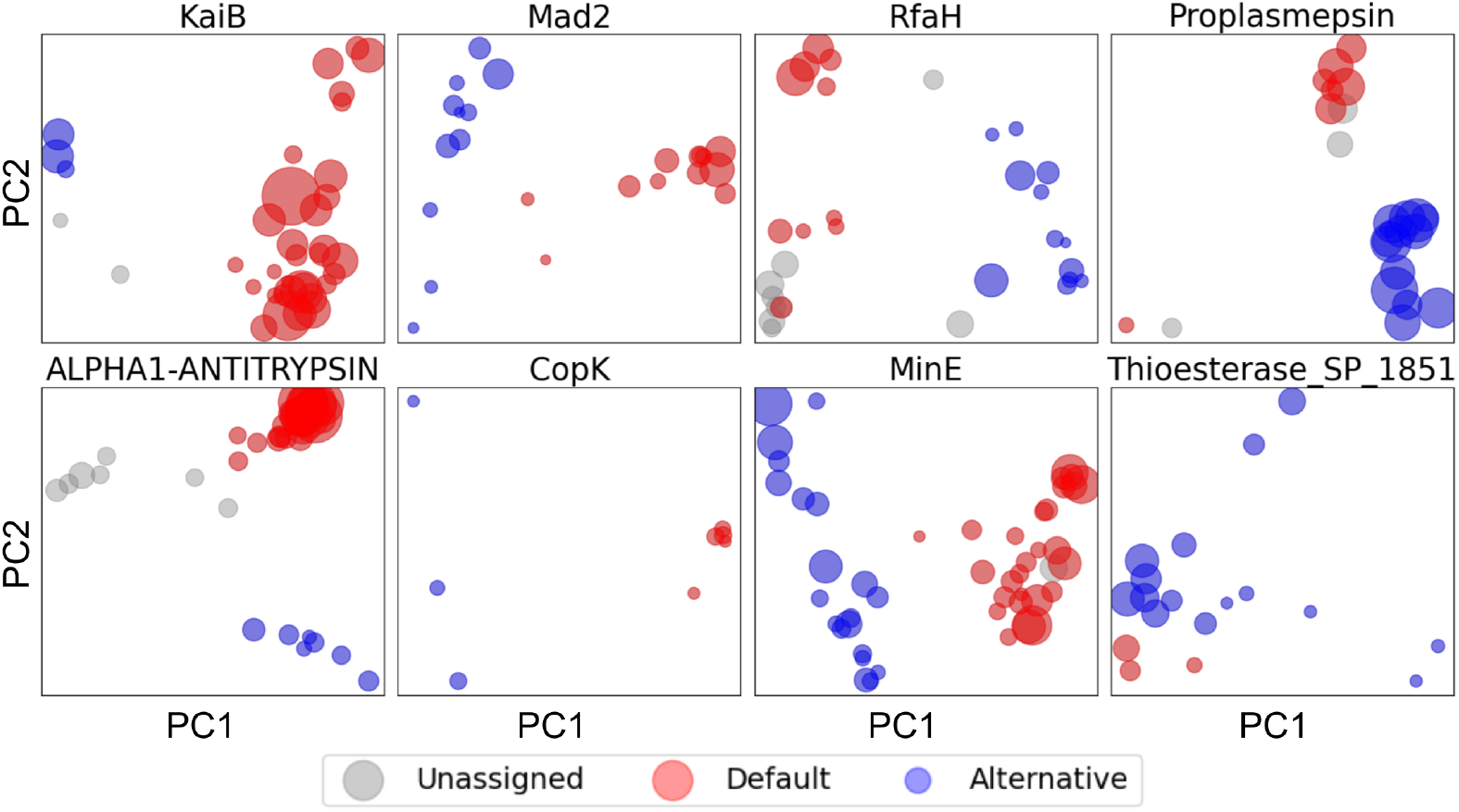
PCA analysis of AF2 Evoformer pair representations of clustered alignments. The size of circles reflects the cluster size.

## 3 Direct Coupling Analysis guides the design of mutations stabilizing conformations

We identify sub-multiple sequence alignments that enable AlphaFold2 to predict the meta-stable state of fold-switching proteins with high accuracy and confidence. The number of grouped sequences is sufficiently large to reliably examine mutation patterns that may influence the predictions, attempting the use of advanced statistical techniques. In particular, we employ Direct Coupling Analysis [11, 12], which focuses on identifying direct evolutionary couplings between pairs of residues by estimating the statistical dependencies between columns in the MSA. It assumes that the residues which are spatially close in the folded structure of a protein tend to co-evolve, meaning that mutations at one residue position often correlate with mutations at another. This co-evolution can be captured by the model and used to predict contacts between residues in the three-dimensional structure of the molecule. It has been shown that extensively sequenced protein families encode co-evolutionary signals corresponding to residue contacts across diverse functional conformational states [14, 20]. Through clustering of the full family, we aim to partition sequences into groups that isolate the covariance signal corresponding to a specific conformation, effectively disentangling it from signatures of alternative contacts. Identifying residue pairs that strongly covary within a particular subset of sequences also serves to prove that the specific subset plays a key role in guiding AlphaFold2 predictions. Moreover, this insight can be leveraged to design targeted mutations that selectively stabilize a particular conformation by introducing substitutions that either disrupt or reinforce these covarying residue pairs. We employ the pydca package [21] using pseudo-likelihood maximization for the inference of couplings [13]. To ensure reliable results from this analysis, sequences should exhibit sufficient variability within clusters. Although DCA is constrained by the limited size of the alignments, their sequences heterogeneity justifies the search for potential co-evolutionary signals (Figure S1 in SI). We then select the top L/2 residue pairs with the strongest couplings from each alternative MSA (L is the sequence length). For each pair, we propose the mutation that shows the greatest divergence in frequency between the alternative and the full alignments. In this way, we aim to identify candidate mutations that promote the stabilization of the alternative conformation over the default one. We first merge the pairs from all clusters and rank them according to their frequency difference, calculated between the clustered and the full MSA. In the upper corner of Figure 6, orange points indicate the top L/2 pairs from our ranked list. In contrast, the red points in the lower corner are the top L/2 ranked couplings obtained from DCA on the full alignment. Contacts, shown in gray, are defined as residue pairs whose C-alpha atoms are less than 12Å apart in the predicted structure. To construct the contact map for alternative states (upper corner of Figure 6), we average residue distances across all predicted alternative structures, while in the lower corner we show the contact map from the default AF2 prediction.

**Figure 6:**
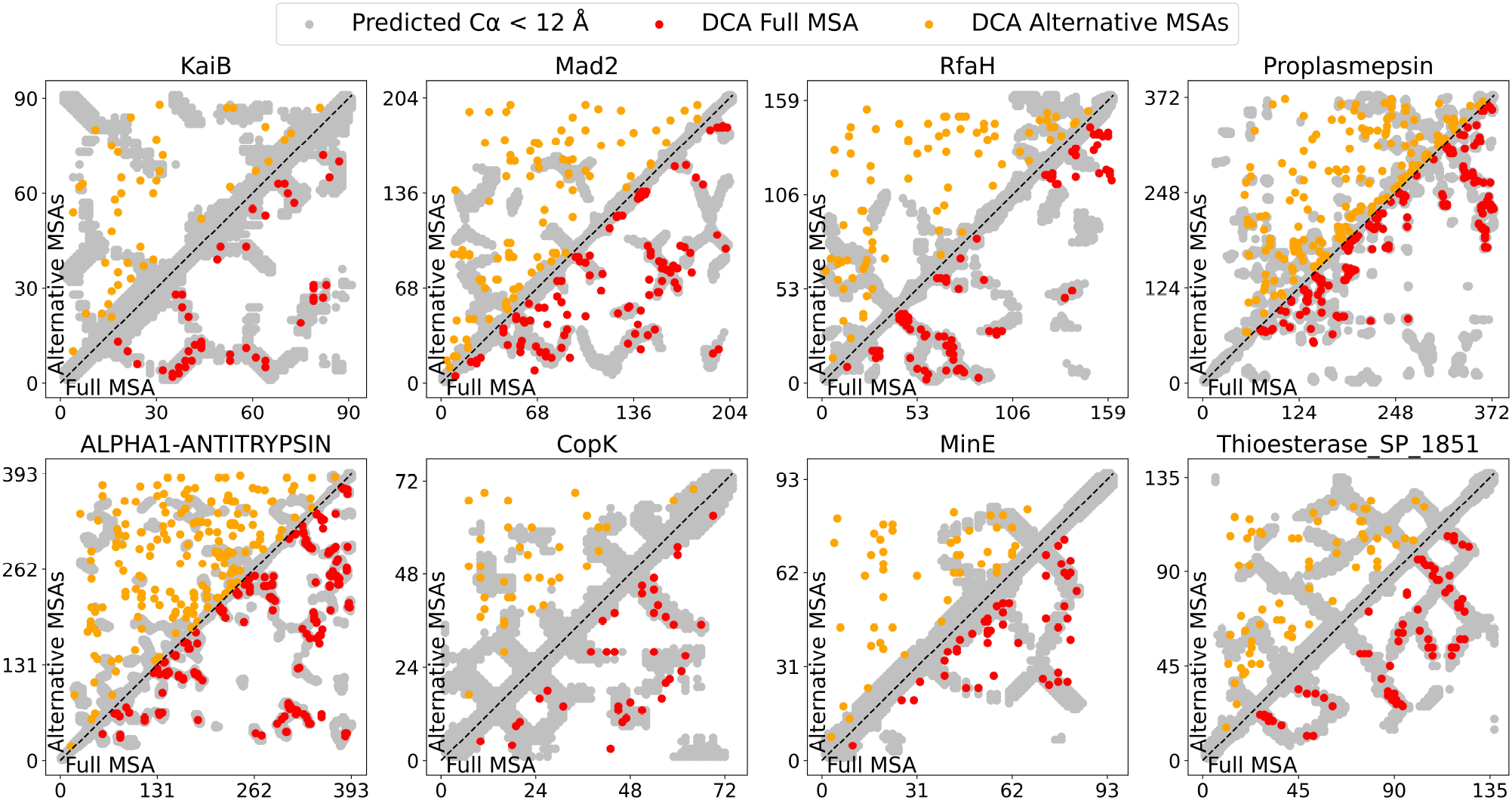
Top L/2 DCA scores from full MSA (red, lower). The top DCA-scored pairs from alternative clusters are merged and ranked by their differential occurrence between the full and clustered MSA, retaining the top L/2 (orange, upper). AF2 predicted contacts with the full and alternative alignments are shown in gray in lower and upper corner, respectively.

The strongest couplings computed from the full MSA align remarkably well with the contact map predicted by AlphaFold2’s default model, confirming that AF2 predictions are biased by the dominant co-evolutionary signals in the protein families, as already discussed in [10]. On the other hand, alternative clusters exhibit a weaker correspondence between couplings and actual residue contacts, as expected due to the significantly smaller alignment sizes and the similarity among clustered sequences. However, several co-varying pairs in the alternative alignments correspond to contacts unique to their respective conformations, and appear crucial in guiding AF2 toward predicting alternative states.

### 3.1 Mutation candidates across proteins

We select from the ranked mutations only those pairs that form contacts in the alternative predicted structure, but not in the default one derived from the full MSA (see Table S1). Among the five mutations identified for KaiB, the double mutation E31H with P67E stands out as a strong candidate for stabilizing the ground state (GS) over the fold-switched state (FS). Specifically, E31H disrupts the salt bridge between E31 and R79 in the folded state, while simultaneously promoting an electrostatic interaction between H31 and E67, which may help stabilize the ground state (Figure S5 in SI). We remind that, for KaiB, the FS corresponds to the AlphaFold2 prediction derived from the full MSA, while the GS refers to the alternative prediction. For Mad2, no mutations are identified using this strategy. On the other hand, seven mutations are proposed for RfaH, with the double mutation P133M-N156L emerging as a key modification. This combination introduces two hydrophobic side chains, which are likely to interact with each other, contrasting with their solvent-exposed conformation observed in the full MSA state (see Figure S6). Proplasmepsin, which displays several large clusters associated with the alternative state, presents 13 mutations. Among them, the combination V59Y-E167K is particularly interesting: the Y-K contact benefits from strong hydrogen bonding and cation–*π* interactions, offering a clear advantage over the hydrophobic-polar mismatch between V and E. Moreover, the E167K mutation may destabilize the GS by disrupting the charge balance with neighboring residues K9, E11, and K16 (Figure S7 in SI). Additionally, the double mutations N61D-R355K introduce a salt bridge in the FS, which could provide stability over the default state. The proposed mutation T50K in CopK would potentially stabilize the alternative state, where K50 and E64 would be close to each other and oppositely charged.

### 3.2 Validation of mutation effect with Molecular Dynamics based Alchemical Free Energy Calculations

Among the candidate mutations discussed above, we selected the KaiB double muation E31H-P67E for validation and use molecular dynamics alchemical free energy calculations (AFEC) to quantify how strongly the mutation favors the GS over the FS. Alchemical methods allow simulation of trajectories in which molecular species are mutated into different ones by switching *on/off* non-bonded interaction of specifically chosen atoms, and the free energy associated with the transformation can be computed by integrating along the alchemical path *λ* (See Figure 7) [15,22]. Early applications of these methods allowed for prediction of the effects of protein mutations on protein stability and drug binding [23], whereas a recent study used AFEC to investigate impact of mutations on protein-protein interface between two signaling proteins. [24]. We aim to adapt these methods to assess how a mutation affects the relative stability of two fold-switching conformations. This is achieved by performing AFEC on both conformers and calculating the difference in Δ*G* values associated with the alchemical transformations, as illustrated in the thermodynamic cycle (Figure 7b). To this end, we adapted the AFEC procedure developed by [25], which employs the Hamiltonian replica exchange scheme (further details in S1). For the E31H-P67E mutation, the resulting ΔΔ*G* indicates a stabilization of the GS conformation over FS by approximately 3.5 kcal/mol, supporting the effectiveness of our pipeline in designing double mutations that influence the relative stability of fold-switching protein conformers. This result further reinforces the hypothesis that co-evolutionary signals play a role in guiding AlphaFold2’s predictions toward specific conformational states.

**Figure 7:**
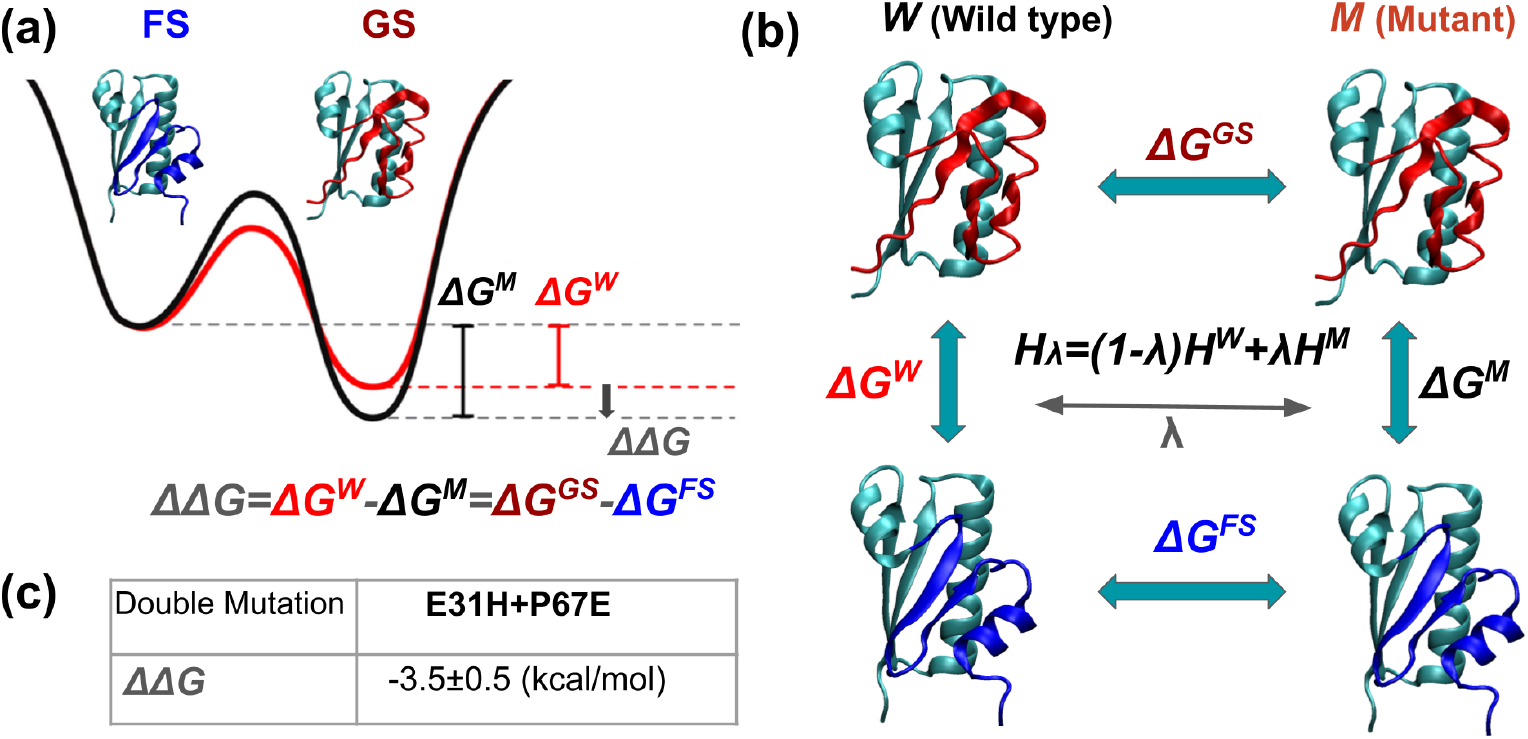
Alchemical free energy calculations assess the impact of mutations on the relative thermodynamic stability of KaiB GS and FS conformers (a), defined as ΔΔ*G* = Δ*G*^*W*^ *−* Δ*G*^*M*^. ΔΔ*G* can be computed as the difference between the horizontal Δ*G* values in the thermodynamic cycle (b), which are computed with MD by integrating along an alchemical path describing the transformation of the wild-type protein into the mutant. (c) The double mutation E31H-P67E in KaiB stabilizes the GS over FS by 3.5 kcal/mol.

## 4 Extending conformation prediction beyond fold-switching

We showcase the versatility of our clustering approach by applying it to proteins undergoing diverse conformational changes beyond fold switching. Specifically, we demonstrate its effectiveness across two distinct protein families: a kinase and a G-protein-coupled receptor. In both cases, we establish a clear correspondence between statistical enrichment and structural outcomes, highlighting the method’s ability to uncover biologically meaningful sequence patterns that shape functional conformational transitions. Consistent with previous findings using MSA sub-sampling [7], our clustering approach identifies both open and closed conformations of the Beta-1 adrenergic receptor, characterized by a slight shift of helix 7 relative to helix 6. However, the predictions reveal more pronounced structural differences beyond these subtle shifts. In particular, one cluster exhibits a significant divergence from the full MSA prediction (Figure 8a), leading to a major fold switch on the extra-cellular side. This transition is defined by a reorganization of sulfur bridge interactions, differing from those in the default conformer. Specifically, the alternative conformation disrupts the native C82–C167 and C160–C166 sulfur bridges while establishing a novel C82–C160 bridge (Figure 8b). Crucially, the statistical properties of this cluster provide a strong rationale for the observed structural rearrangement. The sequences associated with the alternative conformation show a strong enrichment of cysteine at position 160, accompanied by a corresponding depletion at position 167 (Figure 8c). This cysteine redistribution within the clustered sequence directly mirrors the rearrangement of disulfide bonds, reinforcing the idea that evolutionary constraints encoded in the MSA drive the observed conformational change.

**Figure 8:**
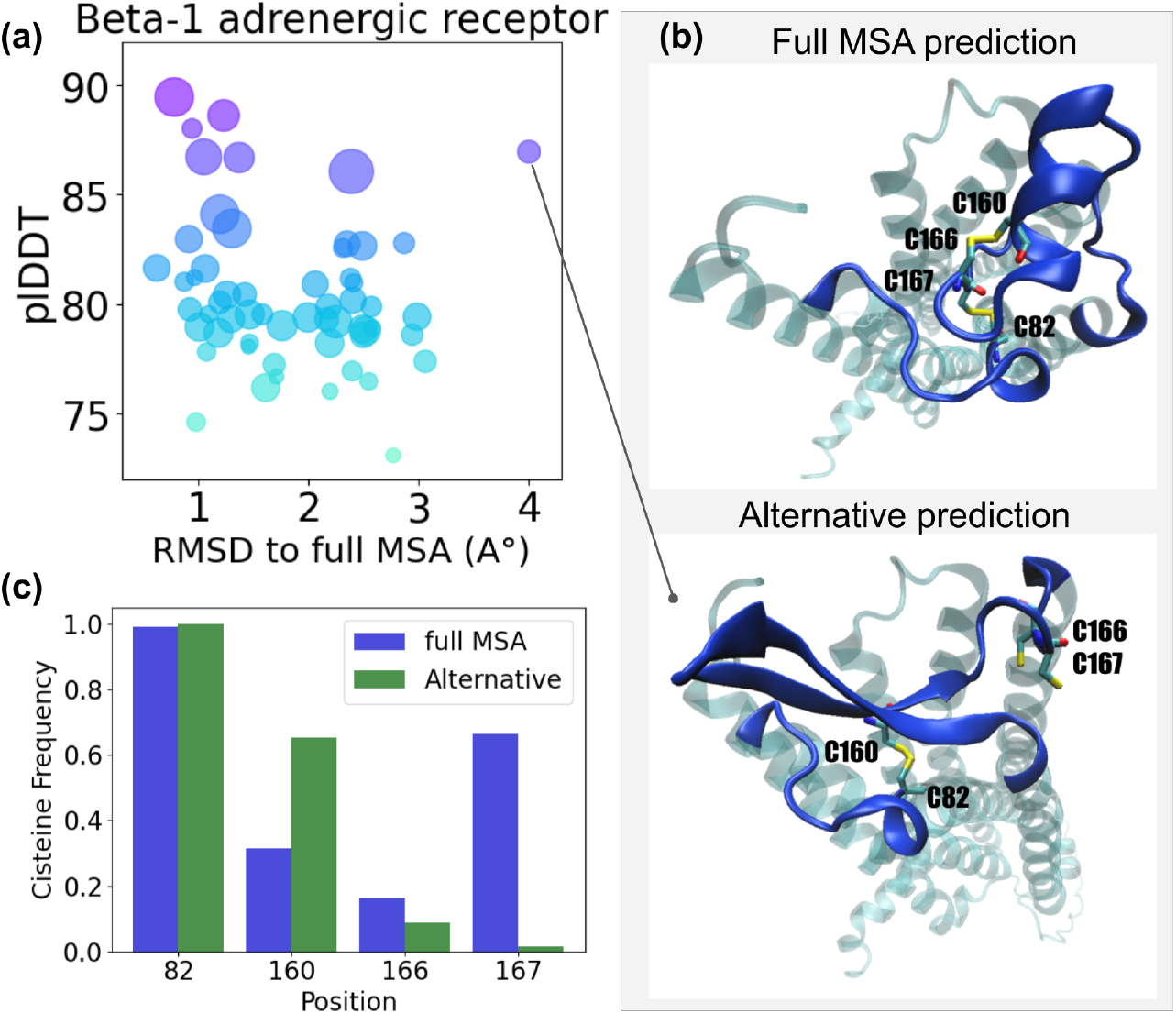
AlphaFold2 structure prediction with clustered MSA for the GPCR Beta-1 adrenergic receptor. (a) Conformation with high plDDT and significant RMSD with respect to the full MSA prediction. (b) Secondary structure rearrangements in the extracellular region: breaking of C82-C167/C160-C166 and forming of C82-C160 sulfur bridges.(c) Cisteine frequencies at rearranged positions in full and alternative MSA.

Building on recent work [26], which demonstrated that subsampling the input MSA of Abl1 Kinase captures both active and inactive conformations, we apply our clustering approach to the Tyrosine Kinase domain of the Epidermal Growth Factor Receptor. Our method successfully distinguishes both functional states by detecting characteristic activation loop rearrangements. Notably, clusters associated with the inactive conformation contain a substantial number of sequences, reflecting a strong evolutionary signal favoring this state. Furthermore, AlphaFold2 predicts the alternative conformation with remarkably high confidence (pLDDT 90%) (Figure 9). Focusing on the MSA linked to the inactive state, we apply our DCA-based mutation detection method and identify a strongly coupled triplet, where position 164 is shared between two residue pairs: 158–164 and 159–164 (Fig. S9). Specifically, position 164 undergoes a K-to-N mutation, reducing positive charge, while residues 158 and 159 mutate from Y to C and H to L, respectively. These mutations likely mitigate the electrostatic repulsion present in the active state, instead stabilizing the inactive conformation, where these residues are spatially close and form favorable contacts.

**Figure 9:**
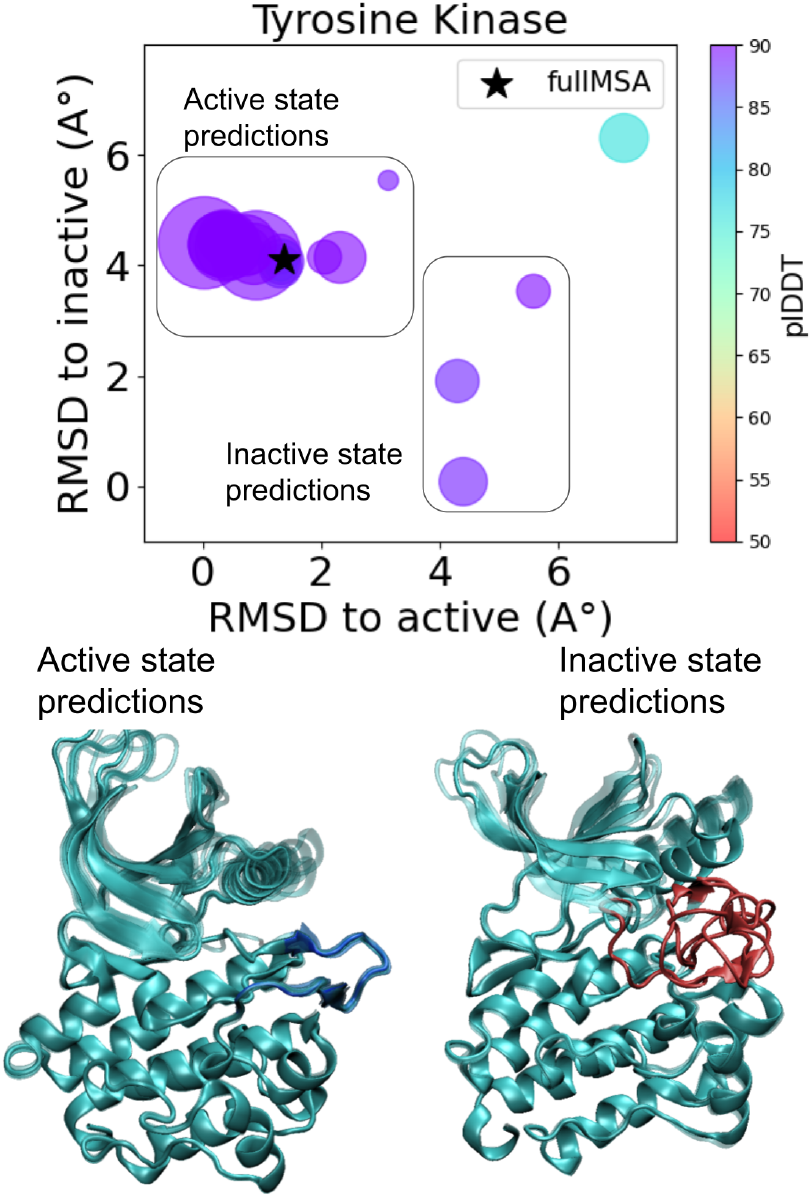
AlphaFold2 structure prediction with clustered MSA for the Tyrosine Kinase: three clusters are associated with the inactive state prediction. RMSDs are computed with respect to AF2 predictions.

## 5 Discussion

We systematically identify metastable states in fold-switching proteins leveraging MSA Transformer-based agglomerative clustering, with greater confidence and efficiency than existing strategies. The effectiveness of the method naturally depends on the availability of homologous sequences. By grouping many sequences, we enable a robust statistical analysis of evolutionary constraints. This allows us to extract meaningful signals linked to conformational changes and design stabilizing mutations using Direct Coupling Analysis, suggesting that AlphaFold2 can be guided by such signals to favor specific conformations. We test the method on eight well-characterized fold-switching proteins, consistently obtaining accurate predictions with high pLDDT scores. The number of false positives from AF2 predictions remains remarkably low, indicating that nearly all clustered sequences yield predictions that closely align with one of the two fold-switching states. Importantly, we are able to identify the alternative state in a protein where the foundational clustering approach failed. Through Principal Component Analysis of the Evoformer module representations derived from clustered alignments, we reveal a clear separation of conformational states at this stage of the AlphaFold2 model. These findings suggest that the MSA representations play a crucial role in refining pairwise representations within the Evoformer, steering the model toward distinct and accurate structural predictions. Furthermore, our clustering framework enables the application of Direct Coupling Analysis within each sequence cluster, allowing us to pinpoint key residue pairs with strong coevolutionary signals specific to each cluster. By leveraging these insights, we can identify residue-residue interactions that are likely crucial in driving conformational transitions between alternative states. The co-evolving pairs identified in the alternative alignments facilitate the design of targeted mutations which exhibit distinct statistical profiles compared to the full MSA, and are predicted to selectively stabilize the corresponding conformational state. To evaluate the impact of the E31H-P67E double mutation in KaiB on protein stability, we perform molecular dynamics simulations. In particular, we use alchemical free energy calculations, a well-established framework for quantifying the effect of mutations on alternative conformations through reliable estimates of ΔΔG values. The results confirm our hypothesis, showing that the E31H-P67E mutation significantly stabilizes the alternative structural state. We further extend the application of our clustering framework to capture a broader range of conformational transitions beyond fold-switching, by applying it to proteins that exhibit distinct structural changes which are functional. We analyze the Beta-1 adrenergic receptor and detect both open and closed states, together with a distinct conformation that diverges significantly from the full MSA prediction, revealing a major fold switch on the extracellular side. The predicted structure disrupts native sulfur bridges while forming a new one, a change that is reflected in the differences in cysteine statistics between the corresponding cluster and full MSA. Similarly, in the Tyrosine Kinase domain of the Epidermal Growth Factor Receptor, our analysis successfully captures characteristic activation loop rearrangements that distinguish the active and inactive states. The mutations identified through our DCA-based approach likely mitigate electrostatic repulsion in the active state, instead stabilizing the inactive conformation where these residues are positioned in close proximity. Our findings reveal key evolutionary patterns within protein families that shape AlphaFold2’s predictions of alternative folds. By refining the clustering strategy, we enhance both the reliability and interpretability of these predictions, as DCA within clusters uncovers meaningful covariant pairs. The detected mutation patterns reflect evolutionary pressures driving the emergence of distinct fold-switching phenotypes, enabling the design of targeted mutations that selectively stabilize a specific structural state. Our results highlight the power of clustering-driven sequence analysis in uncovering biologically relevant metastable states and informing the rational design of mutations to stabilize specific conformations.

## 6 Code and data availability

Code and data are available at https://github.com/RitAreaSciencePark/MSARC.

## 7 Acknowledgments and Funding

The authors acknowledge the AREA Science Park supercomputing platform ORFEO made available for conducting the research reported in this paper and the technical support of the Laboratory of Data Engineering staff. V.P., A.C. and F.C., were supported by the European Union – NextGenerationEU within the project PNRR “PRP@CERIC” IR0000028 - Mission 4 Component 2 Investment 3.1 Action 3.1.1.

## Supporting Information

### S1 Alchemical Free Energy Calculations

In this work, we adapt an AFEC set-up developed in [25], which makes use of the Hamiltonian replica exchange scheme, proposing exchanges every 200 fs. We run 48 replicas simultaneously interpolating bond interactions, Lennard-Jones parameters, and partial charges. We make use of the double system single box strategy to account for the change in net charge induced by the mutations [27]. In particular, we simultaneously simulate the GS and FS KaiB system in the same box, using soft constraints to fix the distance of their centers of mass to 6 nm, and performing the alchemical transformations in opposite directions. In this way, the Δ*G* computed from the single simulation corresponds directly to the ΔΔ*G* assessing the impact fo the mutations on the relative stability of alternative conformers (Figure 7). Simulation boxes consist of rhombic dodecahedrons containing the two proteins, water, Na^+^ and Cl^*−*^ ions with an excess salt concentration of 0.1 M. The systems were energy minimized and subjected to a multi-step equilibration procedure for 8 replica corresponding *λ* values ranging from 0 to 1: 100 ps of thermalization to 300 K in the NVT ensemble is conducted through the stochastic dynamics integrator (i.e., Langevin dynamics) [28], and other 100 ps are run in the NPT ensemble simulations using the Parrinello–Rahman barostat [29]. In production runs, the stochastic dynamics integrator is used in combination with the Parrinello–Rahman barostat to keep the pressure at 1 bar. Equations of motion is integrated with a time-step of 2 fs. Long-range electrostatic interactions are handled by particle-mesh Ewald [30]. Each replica is simulated for 10 ns, for a total of 48 *×* 10 ns = 480 ns. The starting structures for KaiB are selected from the AF2 predictions. To generate a hybrid residue topology, we use the *pmx* packages [31]. Molecular dynamics simulations are performed using GROMACS 2022.5 [32], with the AMBER-ff14SB AMBER force-field for amino acids [33], TIP3 model for water [34], and Joung and Cheatham parameters for ions [35]. Finally, free energy differences are computed with BAR method implemented in GROMACS [36].

**Figure S1:**
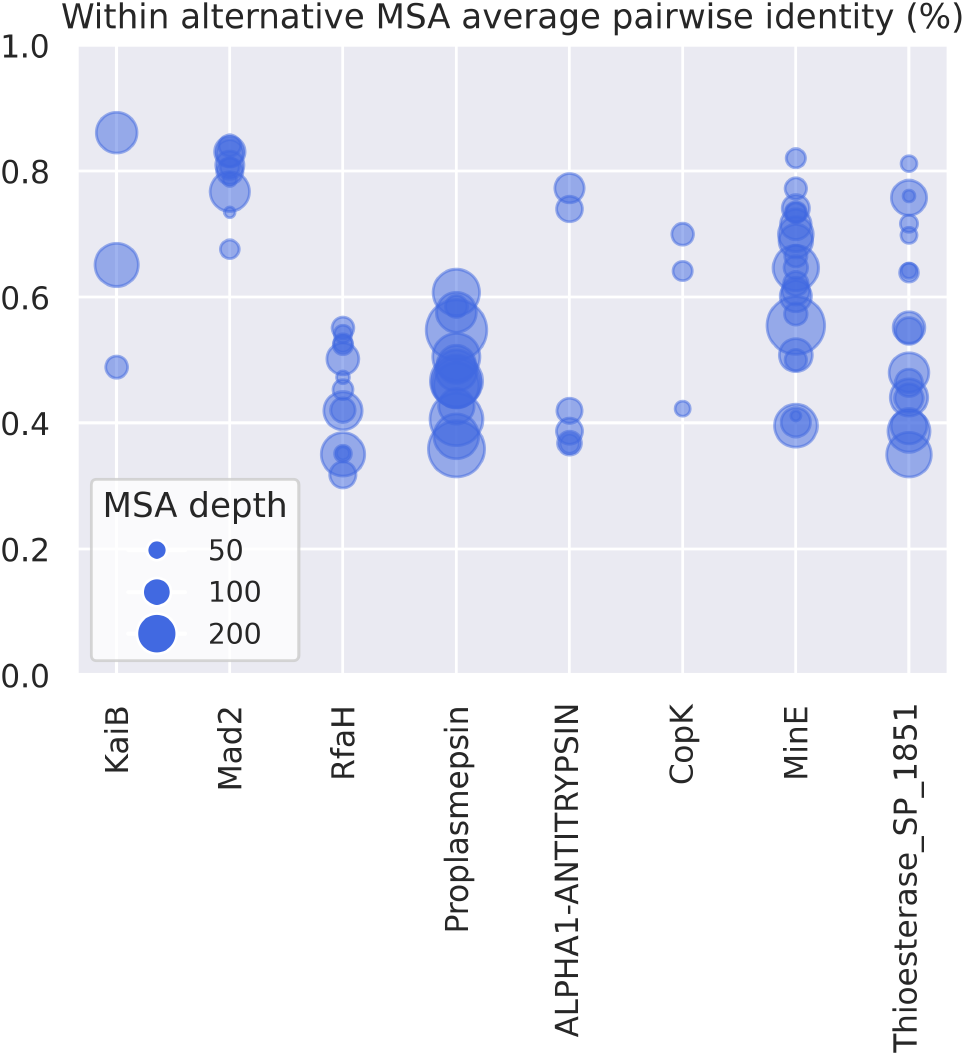
Average % pairwise sequence identities within clusters associated with alternative predictions.

**Figure S2:**
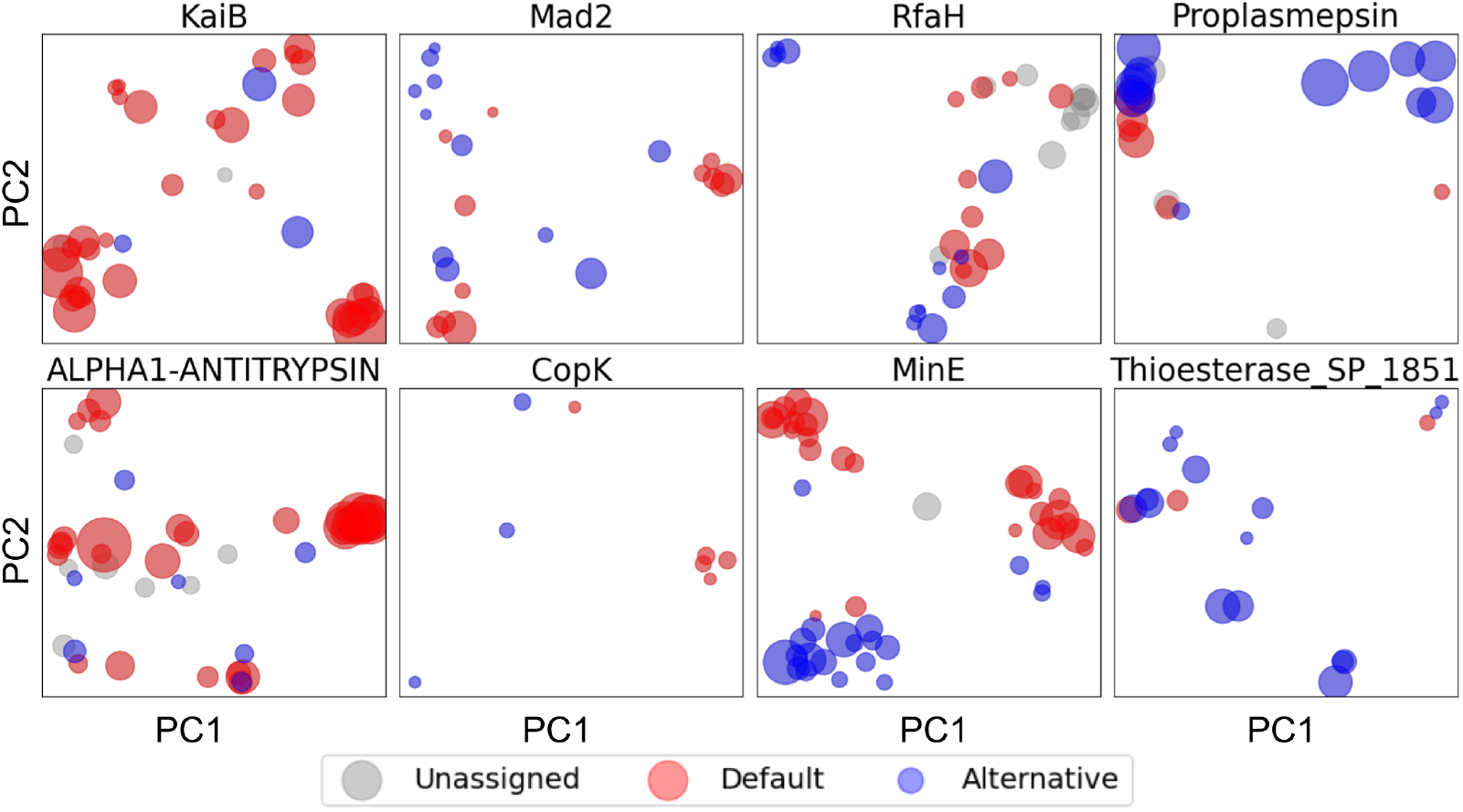
AF2 Evoformer single representations PCA analysis. Red balls correspond to clusters associated to the default AF2 prediction (full MSA state), whereas blue balls are associated to the alternative predictions. Greys balls are unassigned clusters.

**Figure S3:**
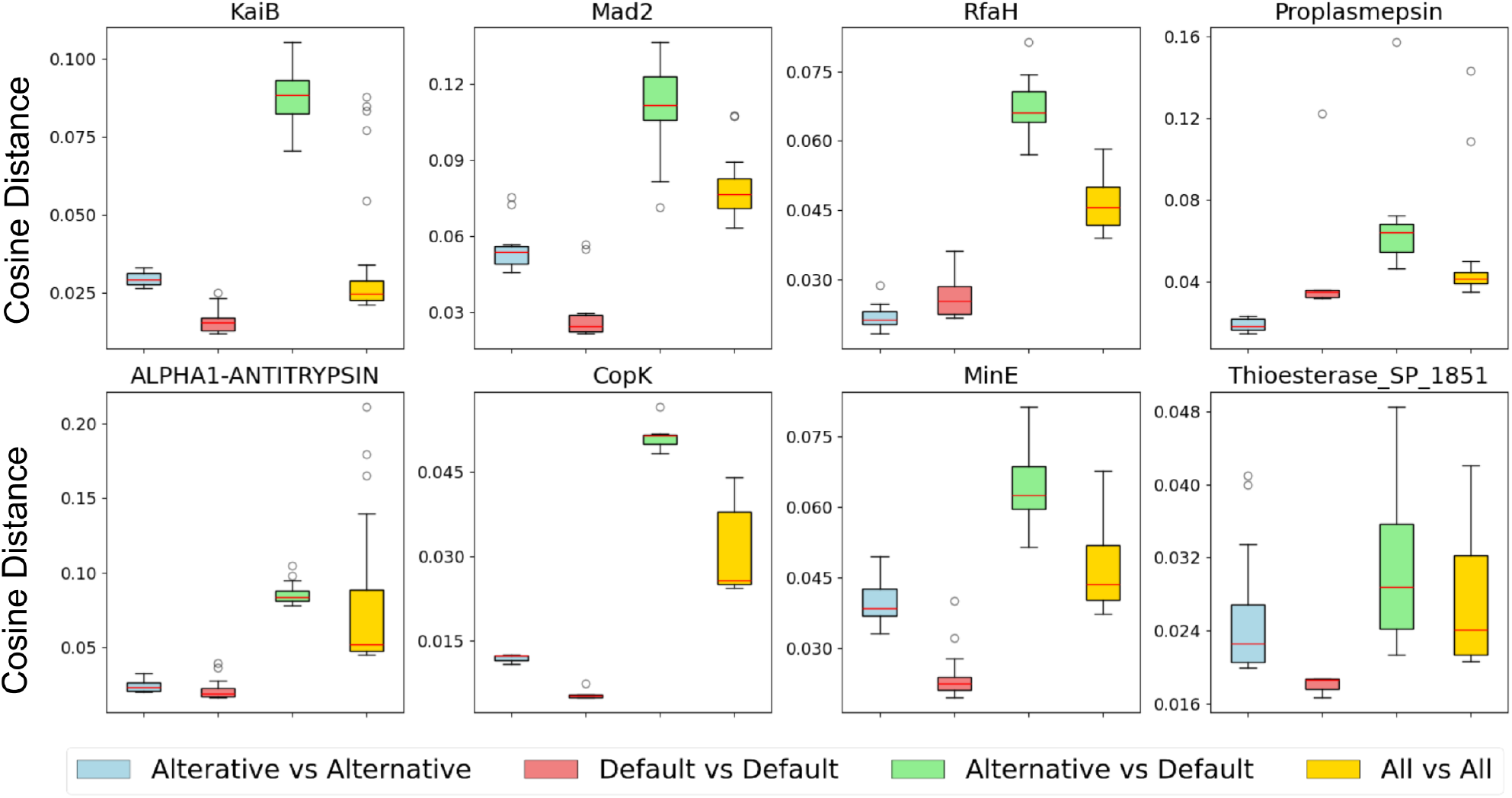
AF2 Evoformer pair representations distances distributions.

**Figure S4:**
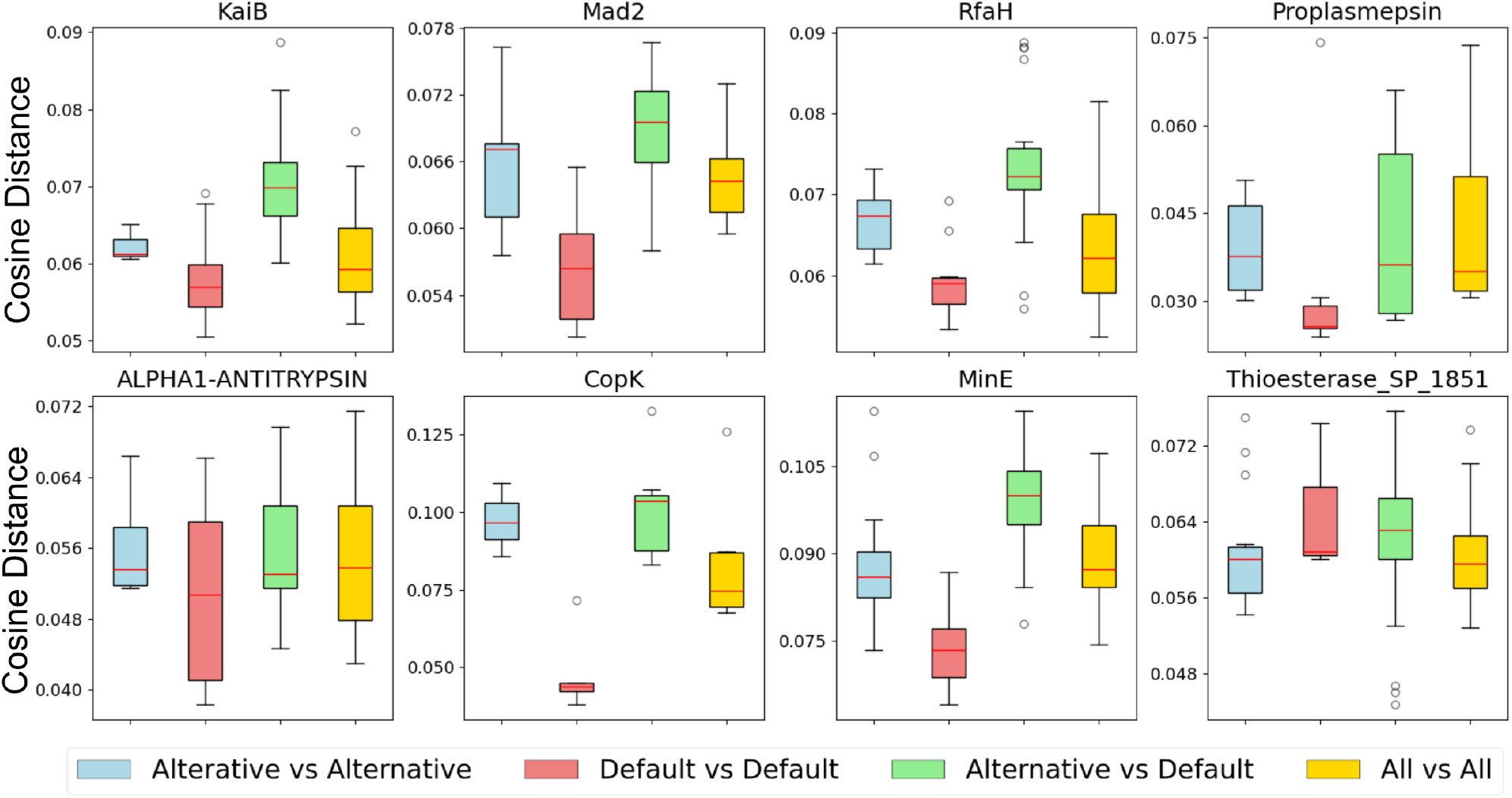
AF2 Evoformer single representations distances distributions.

**Figure S5:**
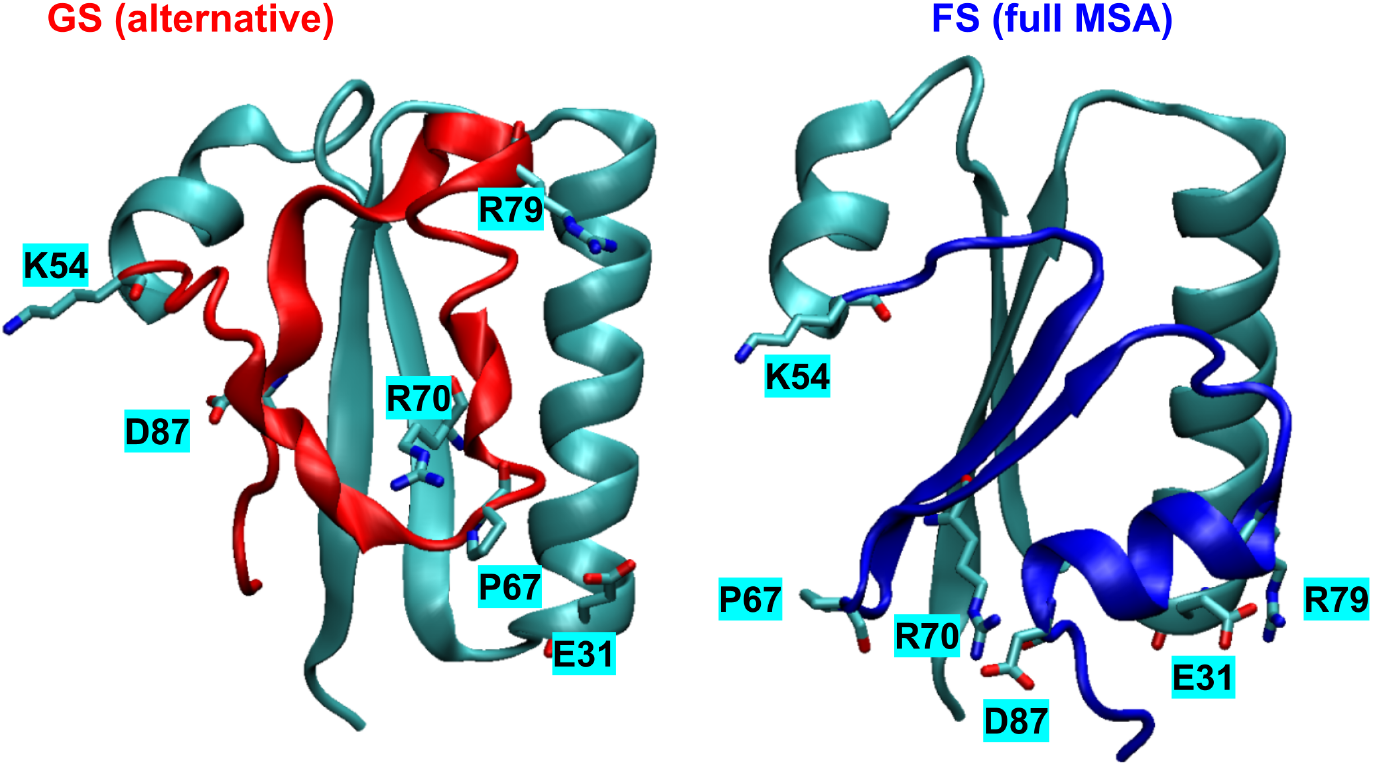

**Figure S6:**
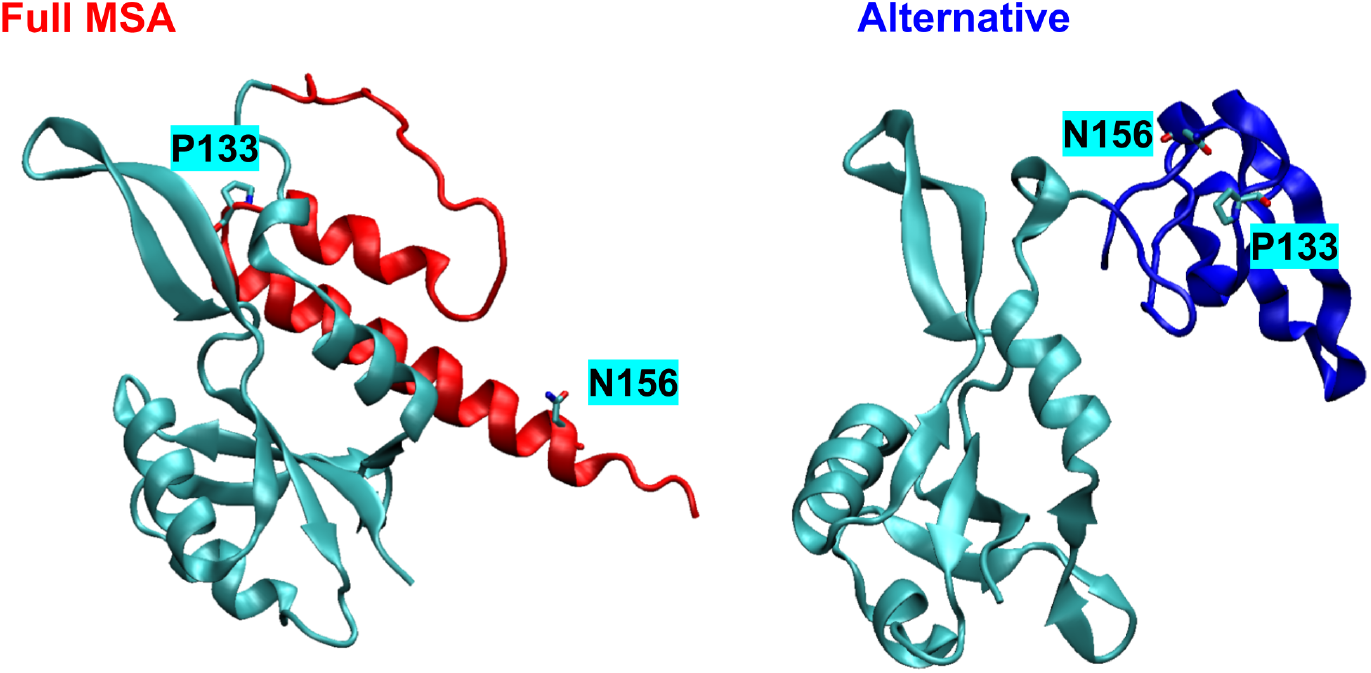

**Figure S7:**
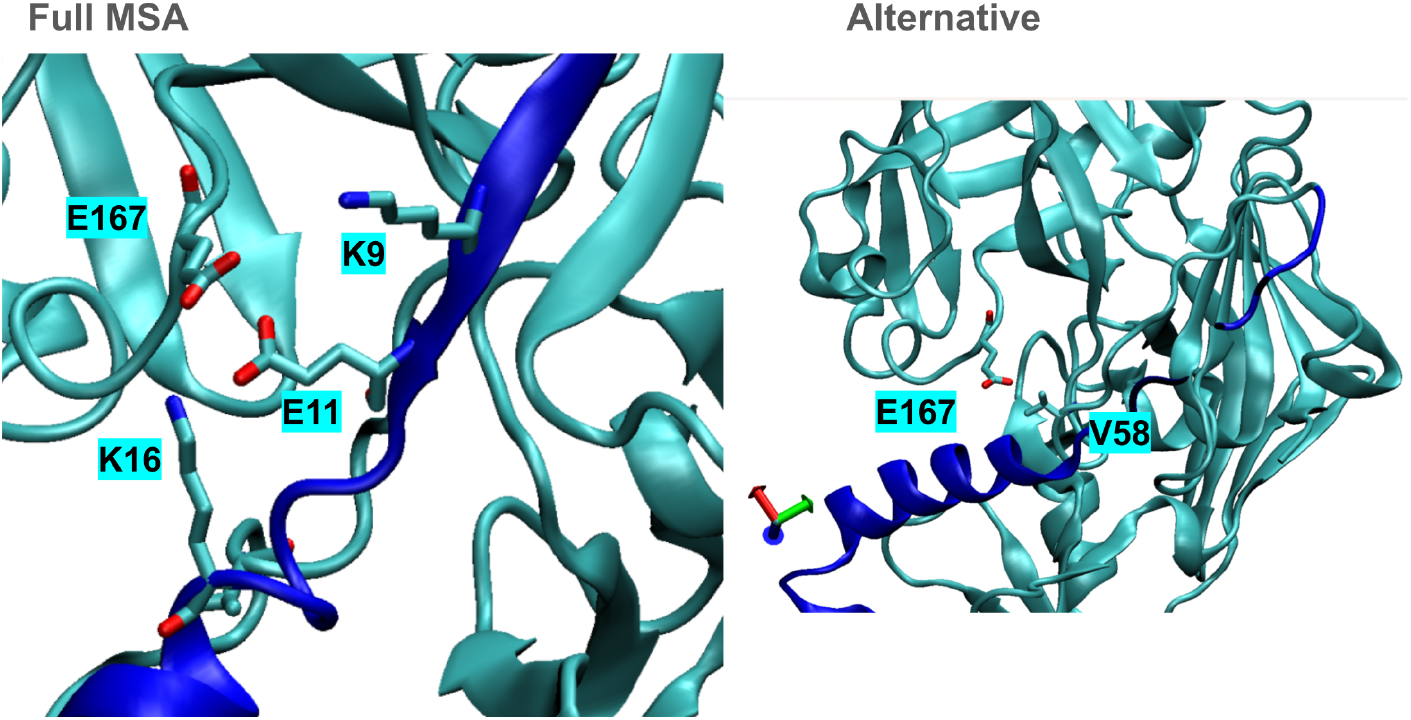

**Figure S8:**
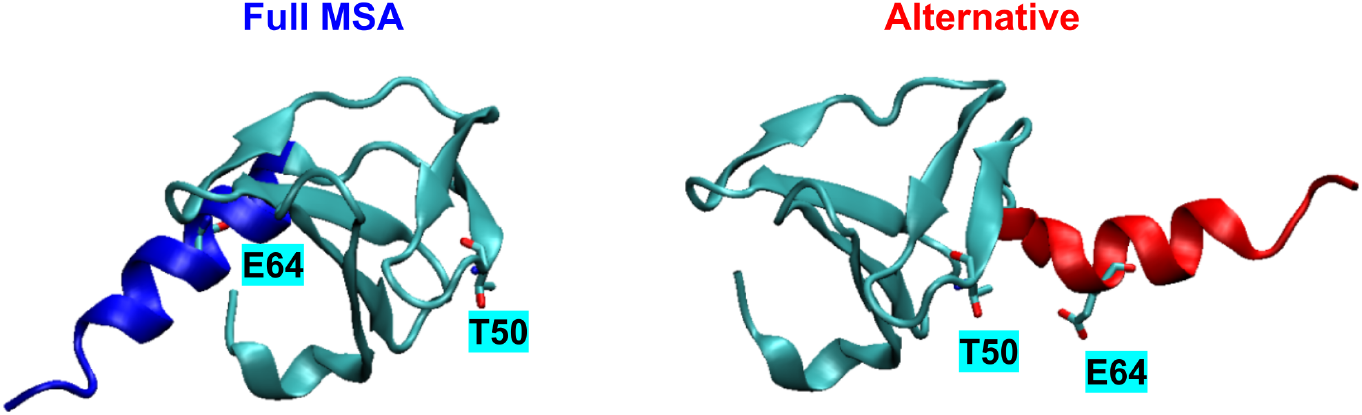

**Figure S9:**
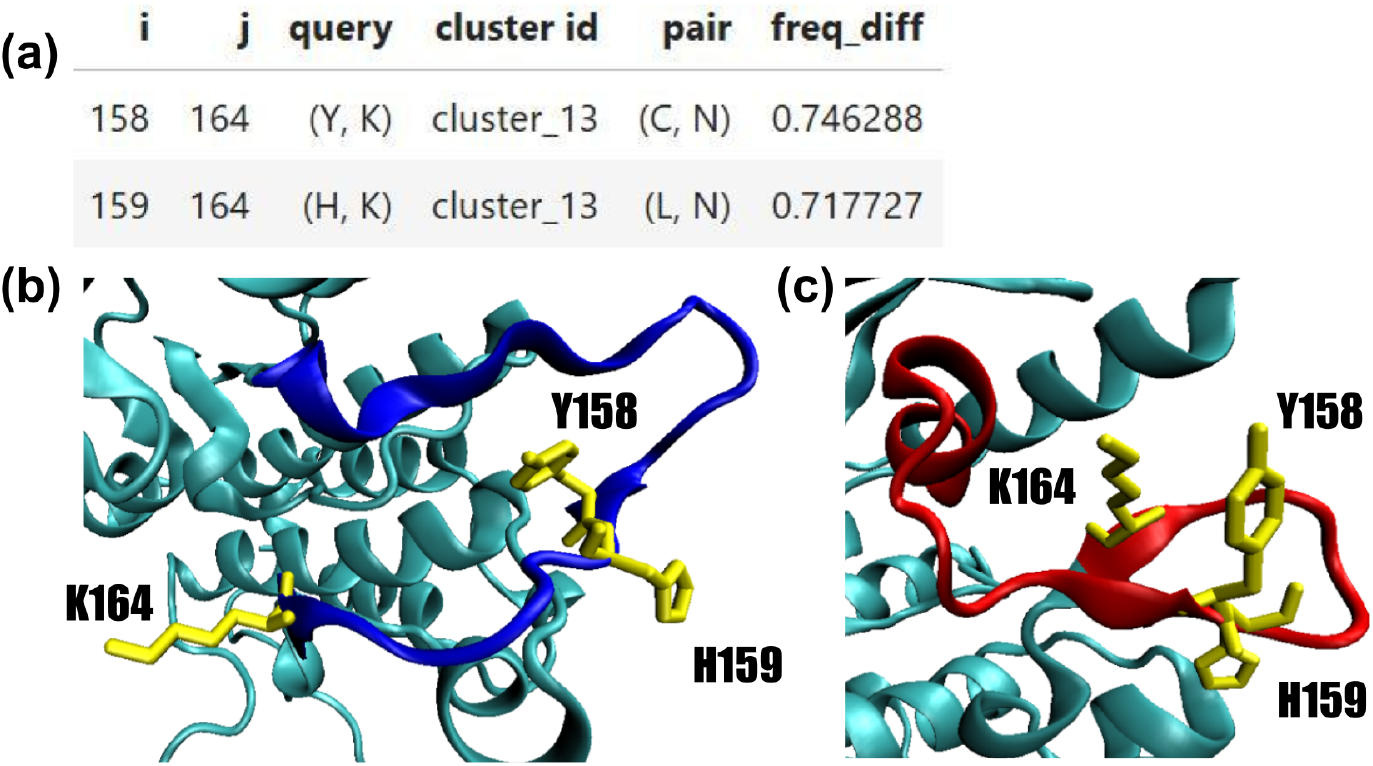
(a) Candidate mutations for Tyrosine Kinase. The shown residues pairs (pair) are highly populated in a cluster (cluster id) leading to the inactive state prediction, with a substantial difference in frequency with respect to the full MSA (freq diff). The triple mutations Y158C-H159L-K164N is supposed to stabilize inactive state (c) over active state (b).

**Table S1:** Suggested mutations for stabilizing alternative state over default.

| System | i | j | Query | Mutation |
| --- | --- | --- | --- | --- |
| **KaiB** |  |  |  |  |
|  | 54 | 87 | K,D | Q,D |
|  | 11 | 80 | A,E | A,D |
|  | 70 | 77 | R,S | K,S |
|  | 31 | 67 | E,P | H,E |
|  | 9 | 77 | Y,S | F,A |
| **RfaH** |  |  |  |  |
|  | 127 | 144 | Q,N | E,T |
|  | 130 | 150 | F,I | Y,V |
|  | 110 | 130 | P,F | P,Y |
|  | 115 | 141 | K,L | R,L |
|  | 133 | 156 | P,N | M,L |
|  | 34 | 99 | T,Y | S,G |
|  | 131 | 154 | T,V | S,L |
| **Proplasmepsin** |  |  |  |  |
|  | 56 | 66 | L,G | I,G |
|  | 58 | 207 | D,P | D,G |
|  | 64 | 207 | F,P | Y,G |
|  | 58 | 213 | D,A | N,G |
|  | 59 | 167 | V,E | Y,K |
|  | 209 | 323 | H,N | G,I |
|  | 59 | 65 | V,Y | V,F |
|  | 60 | 166 | A,V | Q,G |
|  | 69 | 171 | E,I | S,V |
|  | 56 | 213 | L,A | L,G |
|  | 61 | 355 | N,R | D,K |
|  | 24 | 31 | K,K | K,A |
|  | 57 | 80 | D,I | S,V |
| **ALPHA1** |  |  |  |  |
|  | 291 | 345 | L,T | L,V |
|  | 166 | 191 | Q,K | D,D |
| **CopK** |  |  |  |  |
|  | 50 | 64 | T,E | K,E |
| **MinE** |  |  |  |  |
|  | 9 | 14 | G,T | S,T |
|  | 45 | 62 | K,I | Q,I |
|  | 50 | 62 | V,I | I,L |
|  | 11 | 17 | K,V | R,V |
| **Thioesterase_SP_1851** |  |  |  |  |
|  | 13 | 60 | S,L | S,F |
|  | 13 | 60 | S,L | A,Y |
|  | 11 | 16 | A,E | G,L |
|  | 13 | 56 | S,Q | S,S |
|  | 41 | 49 | Y,Y | R,L |

